# Movi 2: Fast and Space-Efficient Queries on Pangenomes

**DOI:** 10.1101/2025.10.16.682873

**Authors:** Mohsen Zakeri, Nathaniel K. Brown, Travis Gagie, Ben Langmead

## Abstract

Space-efficient compressed indexing methods are critical for pangenomics and for avoiding reference bias. In the Movi study, we implemented the move-structure index, highlighting its locality-of-reference and speed. However, Movi had a high memory footprint compared to other compressed indexes. Here we introduce Movi 2 and describe new methods that greatly reduce size and memory footprint of move structure-based indexes. The most compressed version of Movi 2 reduces the Movi index’s space footprint more than fivefold. We also introduce sampling approaches that enable trade-offs between query and space efficiency. To demonstrate, we show that Movi 2 achieves advantageous time and space tradeoffs when applied to large pangenome collections, including both the first and second releases of the Human Pangenome Reference Consortium (HPRC) collection, the latter of which spans over 460 human haplotyes. We show that Movi 2 dominates prior methods on both speed and memory footprint, including both r-index-based and our previous move-structure-based method. The methods we developed for Movi 2 are publicly available at https://github.com/mohsenzakeri/Movi.

## 1. Introduction

Long read sequencing has enabled the building of huge new pangenome collections, such as the Human Pangenome Reference Consortium (HPRC) (Liao et al., 2023) which includes 466 human haplotypes in its recent second release. By using pangenome indexes rather than linear references, we can analyze sequencing reads while reducing the reference bias (Lin et al., 2024; Pritt et al., 2018; Chen et al., 2021). Indexes designed for individual reference genomes are not practical for pangenomes since they scale linearly, i.e. like Θ(*n*) where *n* is the total length of the pangenome collection. Compressed indexes, on the other hand, scale with measures of the non-redundant content in the collection; e.g. the *r*-index (Gagie et al., 2018, 2020) scales like Θ(*r*), where *r* is the number of runs in the Burrows-Wheeler Transform (`BWT`) (Burrows and Wheeler, 1994) of the reference collection. Two compressed index types have proven quite practical: `BWT`-based full-text compressed indexes, (Kuhnle et al., 2020; Nishimoto and Tabei, 2021) and *k*-mer indexes which consist of all substrings of length *k* in the reference (*k*-mers). The size of *k*-mer index grows with the number of distinct *k*-mers in the input, which is related to subword complexity (Kociumaka et al., 2022; Raskhodnikova et al., 2013). Similar to *r*, it reflects the amount of non-redundant sequence in the reference.

The `BWT` reversibly permutes the letters of the input according to the alphabetical order of the letters’ right contexts, i.e. the sequence that comes to each letter’s right. This permutation tends to bring equal characters together into long stretches, where a maximal equal-letter substring of the `BWT` is called a “`BWT` run,” and the total number of runs in the `BWT` is called *r*. When the text becomes more repetitive, the runs become longer and fewer in number. As we add more closely-related genomes to a pangenome, *n* grows linearly while *r* grows more slowly. The ratio *n/r* can be considered the compression ratio achieved.

`BWT`-based indexes are full-text indexes; they allow us to search in the reference for query strings of any length. *k*-mer-based approaches are less flexible and are limited to queries of a fixed length *k*. These approaches are also generally lossy, in that they lose information about continuous sequences longer than *k*. On the other hand, the *k*-mer-based indexes are computationally efficient, making them a practical and widely used method for indexing.

The data structures behind full-text indexes have generally had poor locality-of-reference, and therefore slower query speed. That is, when the index is being queried, the steps of the query process require access to multiple components of the data structure residing in different parts of memory, causing delays while the needed memory is fetched (i.e., it incurs “cache misses”). Recently Movi (Zakeri et al., 2024) was introduced for indexing pangenomes. It is designed based on the move structure (Nishimoto and Tabei, 2021). The move structure is also a BWT-based compressed full-text index, but it has has high locality-of-reference. This means that its memory accesses tend to be predictable and nearby and, as a result, it rarely needs to pause for cache misses. Movi is significantly faster to query compared to other full-text indexes and its query speed is comparable to *k*-mer-based methods.

Similarly to the *r*-index, the move structure grows with the number of runs in the `BWT`, i.e. like Θ(*r*). But the number of bytes required per run (the constant factor) is large compared to other full-text approaches. In this work, we present new ideas to compress the Movi index. The main component of the index is a table with height that grows by *O*(*r*) and width which is a large constant factor. Each row corresponds to a `BWT` run and consists of columns that correspond to the run-length representation, the `LF`-mapping of the run head, and the thresholds information (Bannai et al., 2020) which are offsets in the run. We first introduce a splitting approach that minimizes the space required for the storage of thresholds. We further reduce the size of rows with a scheme that splits rows in order to cap their length. Finally, we present methods for reducing the space required to store the *id* component of the structure, which holds the indexes of the “destination” Movi row containing the `LF`-mapping of the current run’s head. This value needs log_2_(*r*) bits by default, however, we present two approaches for its compression. These ideas exploit the monotonic growth of the values per character, and the relationship between the rows to either sample these values, or use a blocking scheme to encode differences relative to larger components.

Movi 2 is capable of building pangenome indexes that are significantly smaller than Movi’s, while also supporting faster queries. As a demonstration, we measure Movi 2’s index size and speed for three datasets: a set of 94 assembled haplotypes from the HPRCv1 (Liao et al., 2023) release, a larger set of 466 haplotypes from the HPRCv2 release (Liao et al., 2023), and a set of 7,692 complete genomes from 7 bacterial species. We show that the trade-off between index size and query speed is superior using these new approaches compared to either SPUMONI (based on the *r*-index), or to the prior version of Movi. Finally, we explore how Movi 2’s prefetching combines with thread parallelization to achieve further speed gains.

## 2. Methods

### 2.1. Preliminaries

#### 2.1.1. The Burrows Wheeler Transform

The Burrows Wheeler Transform (`BWT`) of *T* is a reversible transformation that permutes *T* ‘s characters according to the lexicographical order of their right contexts (Burrows and Wheeler, 1994). Let *T* be a string of length *n*, |*T* | = *n*, over the alphabet Σ and a special terminal character $ ∉ Σ such that ∀*c* ∈ Σ : $ *< c. T* [*i*] denotes the character at 1-based-offset *i* in T and *T* [*i*.. *n*] denotes a suffix of T starting at position *i. L* = `BWT`(*T*) is a permutation of all the characters in *T* such that for all *i* and *j* such that 1 ≤ *i, j* ≤ *n*, we have *L*[*j*] = *T* [*i*] if and only if the suffix *T* [*i* + 1 .. *n*] is the j^th^ smallest suffix of *T* in lexicographic order. When there are repetitions in *T*, right contexts also repeat, leading to contiguous segments (or “runs”) of identical characters in the `BWT`. The run-length encoded `BWT` (`RLBWT`) compresses a run to a pair consisting of the character and the run length. `RLBWT` is used to build effective compressed indexes for pangenomes (Mäkinen and Navarro, 2005; Mun et al., 2020).

The Burrows Wheeler Matrix (BWM) is a sorted array of *T* ‘s distinct cyclic rotations. The BWM’s last column is BWT(T), as shown in Figure 1.a. The horizontal lack dotted lines in Figure 1.a delineate the `BWT`‘s run boundaries. Let *run*[*i*] denote the *i*^th^ run in the `BWT`, and let *run*[*i*].*h* and *run*[*i*].*t* denote the `BWT` offsets of the first and last letters (respectively) making up the *i*^th^ run. We also call these the run “head” and “tail.” E.g., *B*[7] is the head of *run*[5] in Figure 1.a.

**Fig. 1:**
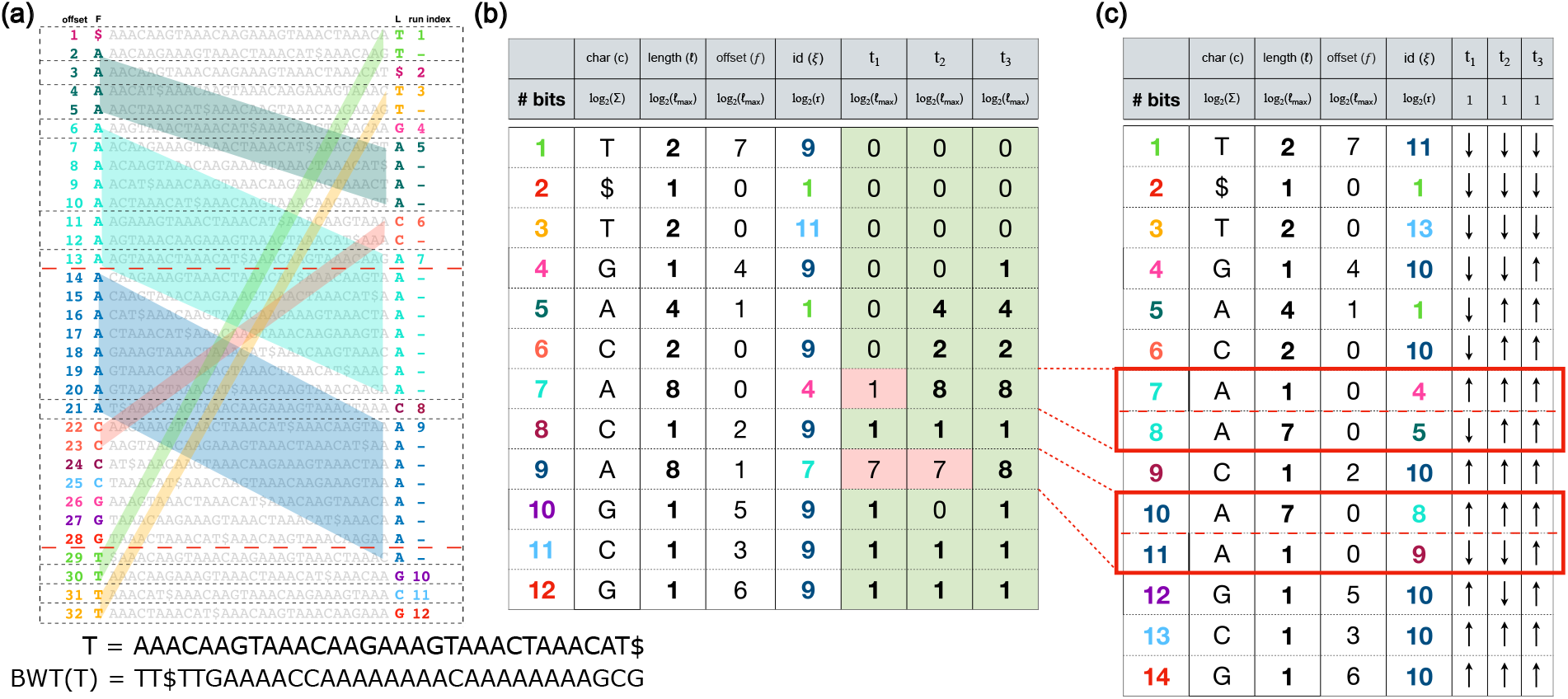
(a) The Burrows Wheeler transform and the Burrows Wheeler matrix built for the sequence T. (b) The Movi index for the sequence T. Non-trivial threshold values have pink backgrounds. Other (trivial) thresholds are either 0 or equal to the length of the run. (c) The Movi table after applying splitting based on the non-trivial thresholds. All the thresholds become trivial, allowing them to be represented with a single up/down bit.

**Fig. 1:**
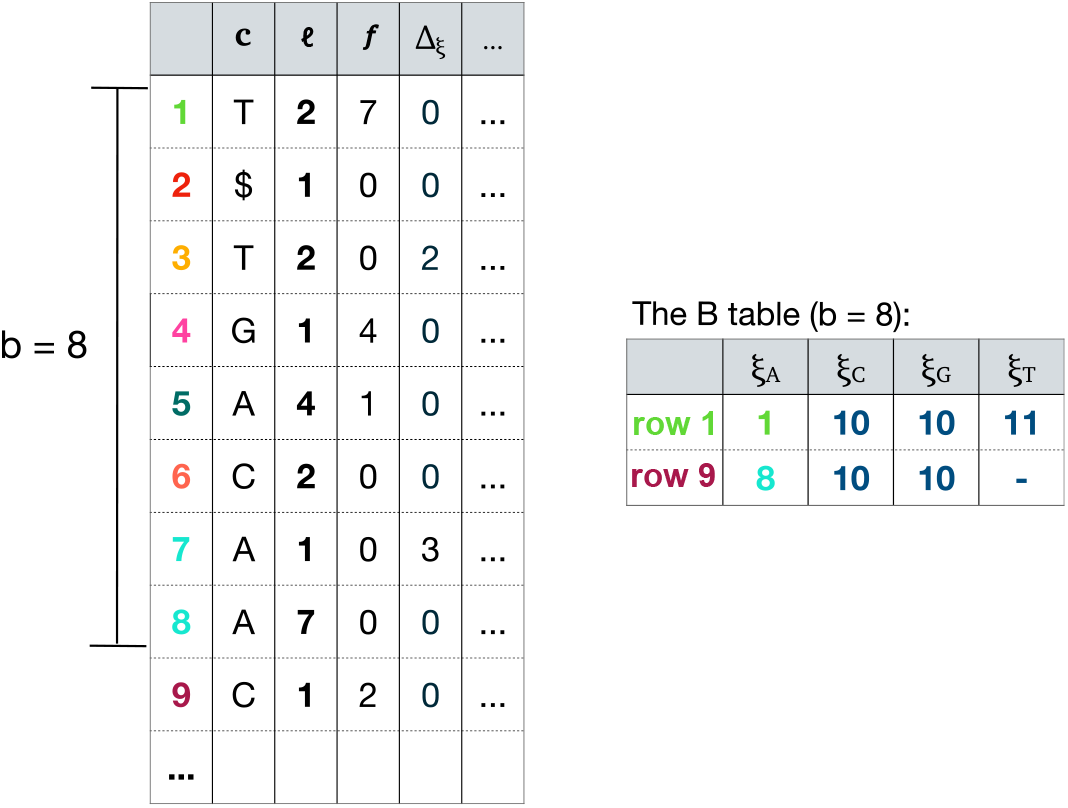
The blocked design of Movi 2 where the exact *id* at checkpoints are stored explicitly for each block in the B table. The block size is 8 in this example. The difference from the checkpoint is stored in the move row.

#### 2.1.2 Thresholds

As detailed later, the algorithm that computes pseudo matching lengths includes repositioning steps where we move several rows either up or down in the BWM. Whether we move up versus down depends on pre-computed values called *thresholds*. For strings *T*_1_ and *T*_2_, the longest Common Prefix (LCP) is the length of the longest string *P* that is a prefix of both *T*_1_ and *T*_2_. For short, we define LCP(*a, b*) as the LCP between rows *a* and *b* of the BWM. Let *run*[*i*] and *run*[*j*] be two runs of the same character (*run*[*i*].*c* = *run*[*j*].*c*) with *i < j* such that no intervening run has the same character, i.e., for every *k* with *i* < *k* < *j, run*[*k*].*c* ≠ *run*[*i*].*c*. The threshold *t*_*i,j*_ is a BWT offset between *run*[*i*].*t* and *run*[*j*].*h* which satisfies these conditions:

1. For all *k* in (run[i].t, *t*_*i,j*_), LCP (*k, run*[*i*].*t*) ≥ LCP (*k, run*[*j*].*h*).
2. For all *k* in [*t*_*i,j*_, *run*[*j*].*h*), LCP (*k, run*[*i*].*t*) ≤ LCP (*k, run*[*j*].*h*).

Any BWM row offset in [*i, j*] that is less than this threshold has a greater or equal LCP with the tail of *run*[*i*] compared to the head of *run*[*j*], and any offset greater than or equal to the threshold has a greater or equal LCP with the head of *run*[*j*] compared to the tail of *run*[*i*] (Bannai et al., 2020). There might several candidates for the threshold between runs *i* and *j* and any one might be selected. Between runs *i* and *j*, multiple candidates for the *t*_*i,j*_ may exist, and any one of them can be chosen. As there are *r* pairs of same-character runs with no intervening runs of the same character, all of the thresholds (one per pair) can be stored in an array of *r* integers.

#### 2.1.3. LF-mapping

The `LF`-mapping property states that the *i*^th^ occurrence of a character *c* in the last column of the BWM corresponds to the same position in the original text as the *i*^th^ occurrence of *c* in the first column (Ferragina and Manzini, 2005). The colored parallelograms in Figure 1a highlight some of these `LF`-mapping relationships. E.g., LF[5] = 32, because both L[32] and F[5] corresponds to position 20 in the text T. Formally, the `LF`-mapping function, *LF* (*i*) = *j*, maps the `BWT` offset i to the offset j where *L*[*i*] and *F* [*j*] are from the same position *k* of the text *T* [*k*]. If *F* [*j*] corresponds to *T* [*k*], *L*[*j*] or BWT[*j*] corresponds to *T* [*k* − 1]. Thus, the `LF`-mapping allows for right-to-left movements with respect to *T*, which is leveraged in pattern-matching queries. The `LF`-mapping function can be computed as:

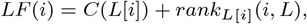

where *C*(*c*) is the number of characters in *T* that are lexicographically smaller than *c*, and *rank*_*c*_(*i, L*) is the number of occurrences of character *C* in *L*[1..*i*] (rank of *c* at position *i* of *L*). For example in Figure 1a, LF[32] = 28 because there are 25 characters (20 *A*s, 4 *C*s, and 1 $) in the text that are lexicographically smaller than *G* and the rank of *G* at offset 32 of the L column (or BWT) is 3.

#### 2.1.4. The move structure

The move structure (Nishimoto and Tabei, 2021) is a compressed full-text index based on the run-length encoded BWTof the text. The size of move structure scales by the number of runs (*r*) in the `BWT`. At the same time, executing an `LF`-mapping movement takes O(1) time. Also of note, the move structure itself consists of a single table. This is in contrast to other compressed full-text indexes like the r-index, which have many components (e.g., bitvectors or wavelet trees), many of which must be queried at each step of the matching process. Because of this difference, queries to the move structure have better locality-of-reference and are substantially faster Zakeri et al. (2024).

There are 5 column for each row in a basic move structure: *c* is the character of the run, *ℓ* is the length of the run, *p* is the `BWT` offset of the run head, *π* is the `LF`-mapping of the run head, and *id* (*ξ*) is the index of the move row (run) that contains `BWT` offset *π*. The parallelograms in Figure 1a illustrate that the `LF`-mapping of a `BWT` offset can be computed based on the distance of `BWT` offset from its run head and the `LF`-mapping of the run head. If *M* is the move structure table, and `BWT` offset i belongs to run index j:

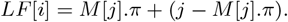

To determine which run index contains *LF* [*i*], we first check whether (*j*–*M* [*j*].*π*) *< M* [*M* [*j*].*ξ*].*ℓ*. If so, then *M* [*j*].*ξ* is the correct next run index. If not, we must scan forward from run *M* [*j*].*ξ* run-by-run until we reach the run containing offset LF[i]. We previously called this “fast-forwarding” (Zakeri et al., 2024). Nishimoto and Tabei (2021) introduced a run-splitting strategy that guarantees a constant number of fast-forward steps while maintaining *O*(*r*) runs. Splitting a run with length *ℓ* will create two new runs with lengths *ℓ*_1_ and *ℓ*_2_ such that *ℓ*_1_ + *ℓ*_2_ = *ℓ*. These runs are no longer maximal and could be termed “sub-runs” or similar.

#### 2.1.5. The Movi index

The Movi index uses a move structure table with *r* rows, prior to any splitting. In the following, we will use “run” to refer to BWT runs which are maximal and “move row” or “row” to refer to rows of the move structure. Note that a move row will often correspond exactly to a BWT run, but can also correspond to a partition of a run, due to splitting. Furthermore, a “row-head” refers to the first `BWT` offset in a move row.

Three of Movi’s columns are identical to those of the move structure (*c, ℓ, ξ*, as seen in Figure 1.b). Unlike the move structure which has columns *p* and *π*, Movi stores only an offset *f*, the distance of *π* from the head of *ξ*^th^ move row. For example, for move row 9 in Figure 1.a,b, the `LF`-mapping of the the row-head is at `BWT` offset 14, which is the second `BWT` offset in move row 7. Its distance from the row-head is 1, so *M* [9].*f* = 1. The strategy for replacing *p* and *π* with *f* was suggested by Brown et al. (2022a).

In Movi, a `BWT` offset is encoded as a pair (*u, v*), where *u* is the index of a move row and *v* is the distance from the head of move row *u*. For example, the last `BWT` offset is encoded as (*r, M* [*r*].*l* − 1). This enables navigation through the text without computing full `BWT` offsets. The `LF`-mapping function in Movi receives a pair (*u, v*) as input and outputs (*u*′, *v*′) representing the `BWT` offset for the `LF`-mapping of (*u, v*). This is computed by first computing an intermediate value, *v*″, the distance of `LF`-mapping result from the head of move row *M* [*u*].*ξ*:

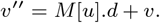

The fast-forwarding procedure determines the move row containing this `BWT` offset and (*u*′, *v*′) is updated according to the cumulative length of the rows skipped during fast-forwarding.

#### 2.1.6 Pseudo Matching Length queries

Pseudo Matching Lengths (PMLs) are an approximation of matching statistics (MSs). MSs form an array of length *m* = |*P* | with entry *MS*[*i*] equal to the length of the longest prefix of the suffix *P* [*i*..*m*] that matches a substring of T. PMLs relax the requirement that *PML*[*i*] equals the maximal prefix of suffix *P* [*i*..*m*] that matches T; rather, PMLs are usually proportional to but less than the MSs. In return for the approximate answer, the algorithm for computing PMLs is faster than those for computing MSs, as shown by Ahmed et al. (2021). Further, PMLs provide similar accuracy in sequence classification tasks compared to MSs (Ahmed et al., 2021).

Computing PMLs starts by comparing the rightmost character of the pattern (*P* [*m*]) to the character at the final offset of the `BWT`, encoded as (*r, M* [*r*].*l* − 1). This process can begin at any offset and we choose the final offset arbitrarily. The process continues by performing iterative `LF`-mapping steps, interleaved with some repositioning steps using the thresholds. The first PML is set to zero, *PML*[*m*] = 0, then for every *k < m* and the `BWT` offset (*i, j*):

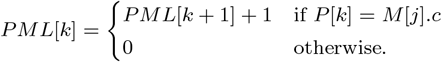

After each *PML*[*k*] computation, in the case of a match (case 1) the `BWT` offset is updated by performing `LF`-mapping. In the case of a mismatch (case 2), we reposition from the current `BWT` offset to a new offset (*i*′, *j*′) such that *M* [*i*′].*c* = *P* [*k*], then perform the `LF`-mapping. Since we always position to a row with a matching character, the `LF`-mapping following the repositioning is guaranteed to succeed.

For repositioning, Movi scans either up from or down from *i*, moving row-by-row until reaching a row with character matching *P* [*k*]. The direction of repositioning is determined by comparing *j* with the threshold value corresponding to *P* [*k*] in *M* [*i*]. If *j* ≤ *M* [*i*].*t*_*P* [*k*]_ then we reposition upward, otherwise downward. Movi must store *σ* − 1 thresholds in a row since a mismatch could involve any character *c*′ ∈ Σ such that *c*′ ?= *M* [*i*].*c*. Note that Movi stores thresholds as an offset in the move row between 0 and *M* [*i*].*l* instead of a global `BWT` offset.

### 2.2. Space efficient indexing with Movi 2

In the default mode of Movi 1, a row of the move table required between 16 bytes to store. For Movi 2, we sought to substantially reduce the space required to represent a single row. Figure 1.b shows the theoretical space required to store each field in a single row. Storing the character (*c*) requires log_2_(|Σ|) bits, and the *id* (*ξ*) requires log_2_(*r*) bits. The length (*ℓ*), offset (*f*) and threshold values (*t*_1_, *t*_2_, *t*_3_) all require log_2_(*ℓ*_max_ + 1) bits, since these are bounded by the maximum length of any move row (*ℓ*_max_).

#### 2.2.1. Splitting by limiting move row length

Some move-structure columns (*ℓ, f, t*_1_, *t*_2_, *t*_3_) require log_2_(*ℓ*_max_ + 1) bits each to store, since their values can range from 0 to *ℓ*_max_. To reduce their size, we set an artificial cap on the length of a move row; i.e. we simply set *ℓ*_max_ according to the number of bits we wish to allocate to these columns, even if some runs exceed that length. We achieve this by splitting any run with length *ℓ > ℓ*_max_ into ⌈*ℓ*_max_*/ℓ*⌉ sub-runs, with all but the last having length *ℓ*_max_ and the last having length equal to the remainder of *ℓ/ℓ*_max_.

#### 2.2.2. Splitting to compress thresholds

In many cases the threshold value for a particular alphabet character, such as *t*_*c*_, is “trivial” in the sense that a mismatch at any of the move row’s offsets should reposition up to the tail of the previous move row with character *c*, or all should reposition down to the head of the next move row with *c*. In this case, the threshold belongs either at the very beginning (offset 0) or the very end (offset *ℓ*) of the move row. For example, in Figure 1.b, many threshold values are set either to 0 or to *ℓ*, indicated by the green background. Only three threshold values are neither 0 nor *ℓ*, indicated by the red background. We call these “non-trivial” thresholds. In Figure 1.a, the red dotted lines highlight the placement of the non-trivial thresholds in the BWM.

If we could eliminate all non-trivial thresholds, a single bit would be sufficient, since it would only need to distinguish whether threshold was at the row’s beginning or end. To achieve this, we propose a new splitting strategy. The goal is to further split the rows of the move structure such that thresholds that were originally non-trivial (i.e. in the middle of a row), such as those denoted with red dotted lines in Figure 1.a), become trivial (i.e. occurring at the move row’s beginning or end). An example of the Movi table after splitting at the thresholds boundaries is shown in Figure 1.c.

The splitting will increase the total number of move rows; however, since there are at most *r* thresholds, at most *r* new move rows would be needed, maintaining an overall size bound of *O*(*r*). In practice, we observe that splitting the runs at thresholds boundaries increases the number of move rows by only *∼*10% of the original number of runs (Table 1). Furthermore, our empirical results demonstrate that the overall effect of thresholds-splitting is a reduction in index size. The thresholds-splitting approach simplifies threshold representation for each character to a binary value, indicating whether the threshold is at the beginning or end of a move row. As a result, storing the thresholds in a Movi index on the DNA alphabet requires 3 bits per move row. This is more than 16 times smaller than the size of the thresholds in Movi 1.

**Table 1.**
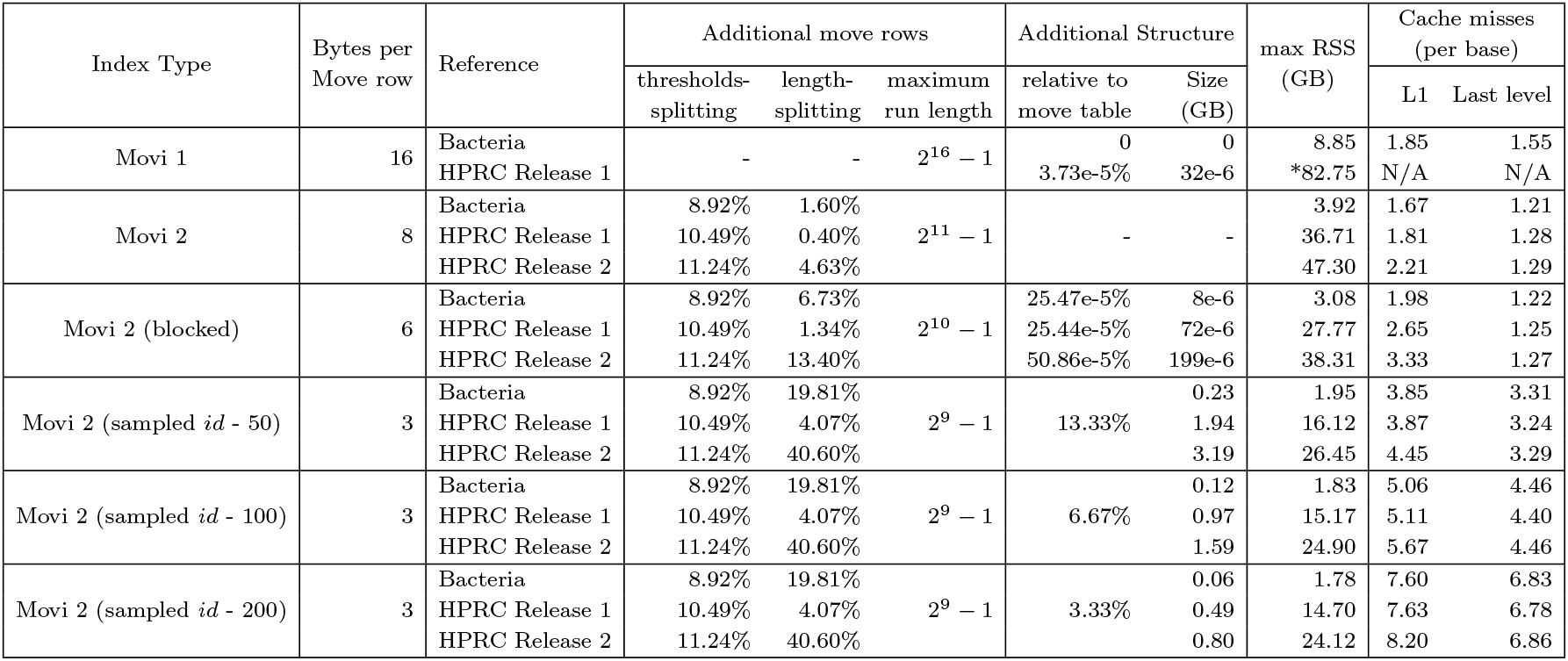
Comparing different characteristics of Movi 2 indexes to Movi 1. Indexes are built on three datasets: Bacteria, HPRC Release 1, and HPRC Release 2. The effect of thresholds and length splitting, the size of the additional data structures, and the number of first and last level cache are shown (cache sizes: L1: 32 KB, LL: 36,608 KB). Movi 1 did not complete successfully on HPRC Release 2 due to high memory usage during index construction. *The Movi 1 index size is larger than the theoretical expectation based on 16-byte row size, because of padding introduced by memory alignment.

In a later version of Movi 1 (1.2.0), another compression approach (discussed in the Supplementary Materials) based on the trivial and non-trivial thresholds was implemented. That approach reduced the size of each move row to 12 bytes by storing at most 1 non-trivial threshold per move row.

#### 2.2.3. Fewer bits to store for the blocked based design

After applying the compression methods described above, the *id* column (*ξ*) will typically become the largest component of each move row, requiring ⌈log(*r*)⌉ bits to store. By default, Movi 2 uses 36 bits to store the *id*, which accounts for more than half the size of a move row. As noted previously (Brown et al., 2022b), an opportunity to compress the *id* field comes from the fact that, for rows labeled by the same character, the *id* field is non-decreasing. In fact, for any symbol *c*, the *id* can be computed by a rank query in a bit vector of size r that has a set bit at the indices corresponding to the move rows with the symbol *c*. This follows from the `LF`-mapping, which stably sorts symbols, ensuring that the *id* of the runs are non-decreasing.

Given the *id* values for all the move rows with the same symbol, one approach could be using a piecewise linear function to estimate the *id* for move row *i* (*id*′ = func(*i, M* [*i*].*c*)) and only store the difference (Δ = |*id* − *id*′|) at the move row. The difference is a smaller value that requires fewer number of bits for storage. This piecewise-linear fitting strategy is inspired by prior work on learned indexing such as the PGM-Index (Ferragina and Vinciguerra, 2020), PLA-index (Abrar and Medvedev, 2024), and Sapling (Kirsche et al., 2021).

In the blocked mode of Movi 2, we divide the move table into equal size blocks of size *b*. Then, we use a step function with fixed length intervals (length *b*) to estimate the *id* for the move rows in each block. For each symbol, the step function returns the *id* of the first move row with that symbol in the block. These values are also called “checkpoints”. The difference (Δ) from the checkpoints are stored in the move row instead of the absolute value of *id* (*ξ*). An example is shown in the Supplementary Materials Figure 1 where *b* is set to 8. The Δ column is stored instead of the *id* column in the move rows.

The *id* at the checkpoints are stored in a separate table called the *B* table. For a checkpoint at *i*^*th*^ move row, we store |Σ| different *id* values, one for each symbol in the alphabet. The stored *id* for a symbol *y* corresponds to the *id* of a move row with the smallest index *i*′ such that *i*′ ≥ *i* and *M* [*i*′].*c* = *y* (i.e., *i*′ is the first occurrence of *y* at or after *i* in the block).

If we choose *b* large enough, the number of blocks is reduced such that the *B* table fits into cache. That makes the retrieval of checkpoints very fast. In practice, we see that the number of last level cache misses never increases by more than 1%, and in some cases, even decreases. For the first level cache, depending on the size of the *B* table, cache misses increase by 19% to 51%. Overall, this approach achieves space efficiency without noticeably compromising speed.

We can use the Δ_*id*_ and the B table to recompute the original *id* values for a row *i* with character *c*. The closest checkpoint before *i* is computed as *i*′ = *i/b*, then the *id* is computed as:

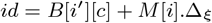

#### 2.2.4. Sampling the *id* field at the checkpoints

We can extend the blocking idea to a sampling approach which does not store Δ values explicitly. Instead, the Δ may be inferred by computing the distance of a row-head from a checkpoint, i.e., head of a sampled move row, by scanning the move rows in between. We denote the sampling rate by s and employ the same method as in the blocked design to store the *id* for each symbol at the checkpoints. In this mode, the table containing the *id*s of sampled move rows is referred to as the S table.

To find the *id* value of a move row *q* with symbol *y*, we examine move rows between *q* and the closest sampled row. Assume the closest sampled move row is at index *i < q*. First, we must scan the move rows [*i, q*) and compute the cumulative length of rows having symbol *y*. This will be the distance in terms of BWT offsets between heads of move rows *i* and *q* after `LF`-mapping. The distance after the `LF`-mapping is important, because the *id* of move row *q* is the index of the move row containing the `LF`-mapping of the head of move row *q*. We can now compute the *id* of move row *q*, given the *id* for symbol *y* at sampled move row *i* and the corresponding offset for that. The offset of the sampled move rows can be retrieved by simply examining the content of the move row, since *f* (the offset) is stored for all rows.

The process at this point, becomes similar to a fast-forward step. Given the `LF`-mapping of a BWT offset, we want to compute the `LF`-mapping of another BWT offset at a certain distance. This step invokes another scan from the retrieved *id* for *y* until the desired BWT offset is reached. We illustrate this procedure through an example in Figure 2 where we compute the *id* of row 7 based on the sampled move rows. A pseudo code of this process is also provided in Supplementary Materials Algorithm 1.

**Fig. 2:**
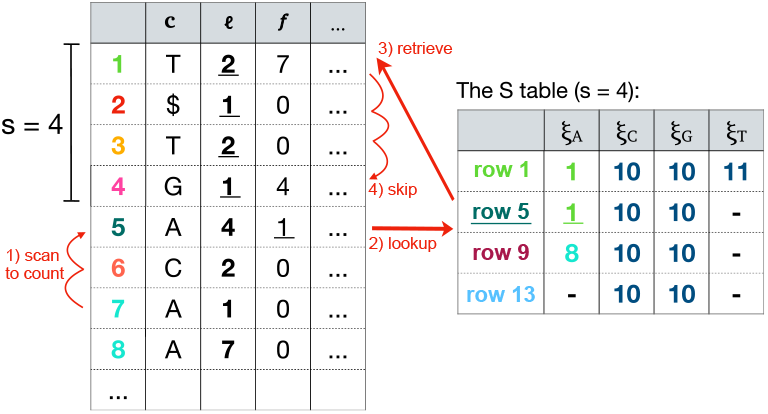
The sampled design where no *id* is stored in the move rows. Instead, *id* at sampled move rows are stored in the S table. As an example, the *id* of move row 7 is computed in four steps: 1) scan to count the 4 As between the move rows 7 and sampled row 5, 2) lookup the *id* for A at row 5 which is 1, 3) retrieve the move row identified by the *id* (row 1), and 4) scan move rows until 4 BWT offsets (as many as the number of As counted in step 1) are skipped from the BWT offset (1, 1). The final move row reached (row 4) is the *id* of move row 7.

## 3. Results

### 3.1. Analysing different versions of Movi 2

To compare different Movi 2’s indexes to Movi 1, we used three different datasets: (1) a collection of of 7,692 complete genomes from 7 bacteria species (“Bacteria”), (2) 94 human haplotypes from the first release of Human Pangenome Reference Consortium (“HPRC Release 1”) (Liao et al., 2023), and (3) 464 human haplotypes from the second HPRC release (May 12, 2025) supplemented with CHM13 (Nurk et al., 2022) and GRCh38 (Schneider et al., 2017) (“HPRC Release 2”).

First we compared different versions of the Movi 2 index with Movi 1 and evaluated how Movi 2 compression strategies affected different aspects of the index. Movi 1 uses 16 bytes per row, while the default Movi 2 uses 8 bytes. The blocked design uses 6 bytes and the smallest version (sampled-*id*) uses 3 bytes per row. The sampled-*id* version provides speed-memory trade-off by varying the sampling rate. Here we tried 50, 100, and 200.

The smaller row size in default Movi 2 is achieved via two run-splitting strategies. The first, thresholds-splitting, forces all the thresholds to be trivial, allowing a single threshold status bit to be stored instead of a full threshold value. The second, length splitting, reduces the bits required to store length and offset columns. As there are at most r distinct thresholds, in the worst case, the thresholds-splitting doubles the number of rows. However, in practice we observed far fewer rows are added. Table 1 shows that thresholds splitting increased the number of move structure rows by 8.92% to 11.24% in different datasets.

Table 1 also shows the percentage increase in rows due to length-splitting. In default Movi 2, we use 11 bits for the length (*ℓ*) and offset (*f*) columns. For the blocked index, we use 10, and sampled-*id* uses 9 bits. When fewer bits are used, more splitting is required. With maximum run length of 2^11^ −1, the number of rows was increased by 1.60%, 0.40%, and 4.63% for the Bacteria, HPRC Release 1, and HPRC Release 2 indexes respectively. For the blocked indexes, the increase in the number of rows was between 1.34% to 13.4%. The largest growth in our experiments is seen for the sampled-*id* variants, where the number rows was increased between 4.07% to 40.60%.

For length-splitting, we observed that the amount of splitting required was dataset dependent. Highly repetitive datasets will also have high *n/r* and so (by definition) longer average run lengths. The Bacteria dataset spans many species but includes some collections of thousands of genomes from the same species with *n/r* (average run length) of 157.94. The HPRC Release 1 dataset, consists of 94 human haplotypes and has an *n/r* of 133.80 while the HPRC Release 2 includes 466 human haplotypes with *n/r* of 535.02. For HPRC Release 2 dataset, we saw an increase of 40.60% in the number of move structure rows when the maximum run length is bounded by 2^9^ − 1. However, since the sampled-*id* version only uses 3 bytes per row (more than 80% less than the original Movi index), the overall size of the index becomes smaller. The “Additional structure” column in Table 1 shows the absolute and relative size of the additional table used in some indexes compared to the main move structure table. For Movi 1, this additional table stores rare move rows that do not fit in the main table because their length, offset, or threshold values require more than 16 bits to represent.

For the blocked and sampled-*id* indexes, this column shows the size of the B table and S table respectively. The size B table depends on the number of checkpoints and the size of the blocks. It was small in our experiments: 8 KB for Bacteria, 72 KB for HPRC Release 1, and 199 KB for HPRC Release 2. For HPRC Release 2, we had to select a smaller block size to guarantee all deltas could fit in the delta-*id* column. For the sampled-*id* version, we evaluated sampling rates of 50, 100, and 200, which result in additional tables that are 13.33%, 6.67%, and 3.33% of the size of the main table, respectively.

To assess how the compression strategies affected locality of reference, we used the “Cachegrind” profiler to measure cache misses at the first (L1) and last (LL) cache levels. Default Movi 2 had lower L1 and LL (last level) cache misses due to its smaller size compared to Movi 1. For the variants with extra table (blocked and sampled-*id*), the number of cache misses were increased due to extra access to either the B or S table. The B table is small enough to fit in the Last Level cache of most systems. This was evident as the number of LL cache misses were not significantly increased, and even decreased in some cases due to smaller size of the main table. For the sampled-*id* index, the number of cache misses increased with higher sampling rates because more scanning in the main table was needed to get to a sampled move row.

### 3.2. Movi 2’s space/speed trade-off

We performed benchmarks to assess the space/time trades achievable by Movi 2’s compression methods. We used long reads to compare to compare Movi 2’s PML query to the same query performed by Movi 1 and SPUMONI (which uses the *r*-index). We also included ropebwt3 (Li, 2024) in our benchmark which is another scalable tool for full-text indexing of pangenomes, though it supports a different and broader set of queries compared to the other tools in the benchmark. In this experiment, ropebwt3 was used to compute Super Maximal Exact Matches (SMEMs).

For the bacteria dataset, we used 100K ONT reads from SRR11071395, the Zymo High Molecular Weight Mock Microbial Community (Kovaka et al., 2021). These had an average length of 15K bases. For the HPRC datasets we used PBSIM2 (Ono et al., 2021) to simulate 1.18 M reads from the CHM13 with an average length of 2.6K bases.

Figure 3 presents the results, with five points corresponding to the different compression/speed trade-offs implemented in Movi 2, alongside the results from Movi 1, SPUMONI, and ropebwt3. The fastest method was default Movi 2 which also had a lower peak memory footprint compared to Movi 1. PML query with this mode was about 35 times faster than the same query performed by SPUMONI. The default Movi 2’s index was 56% smaller than Movi 1 in all three different datasets. The PML query with this Movi 2 index was also between 27% to 30% faster than Movi 1, likely due to its decreased rate of cache misses and smaller index. The next fastest mode of Movi 2 is the blocked mode which was between 19% to 24% smaller than default Movi 2. This mode required 65% less memory compared to Movi 1. Query with the blocked Movi 2 was about 13% faster than Movi 1 for the HPRC Release 1 dataset. For the bacteria dataset, the speed of blocked mode query was less than 1% slower than Movi 1.

**Fig. 3:**
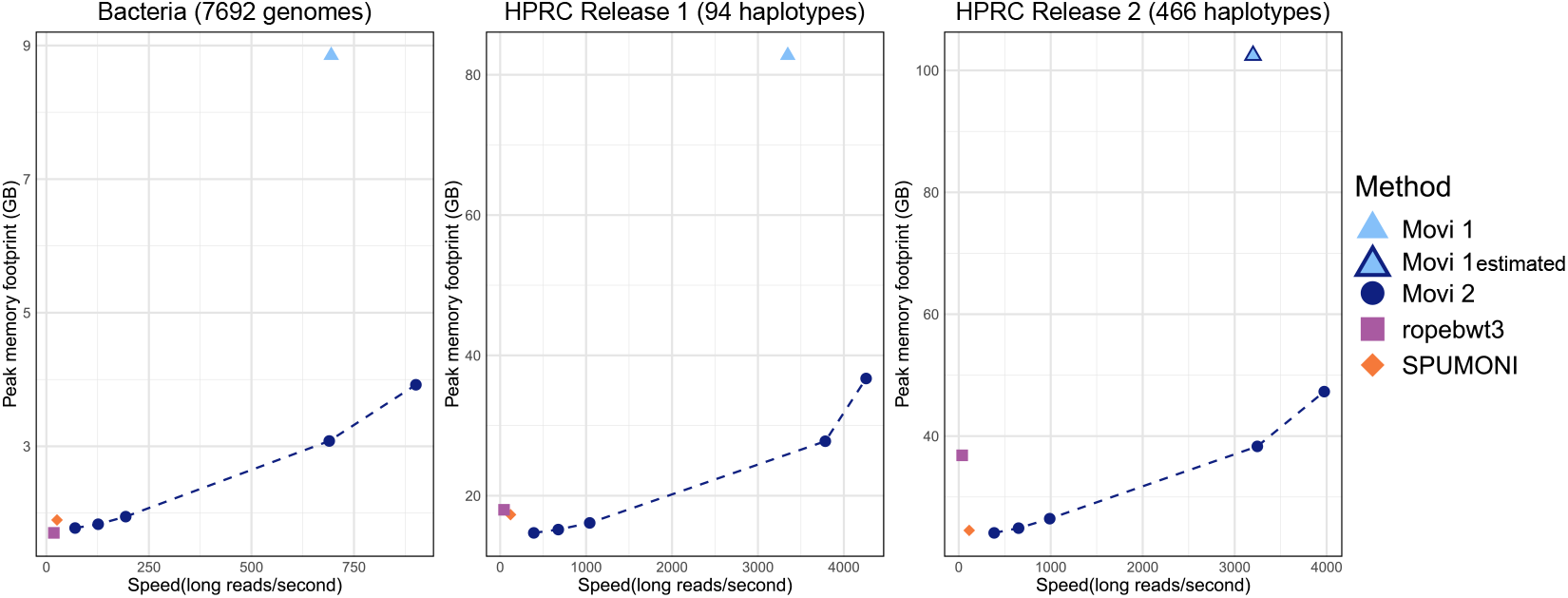
The time and maximum resident set size (max RSS) for querying different indexes. Movi, Movi 2, and SPUMONI compute PMLs while ropebwt3 computes Super Maximal Exact Matches longer than 19. The average of 5 consecutive runs for each method is reported. The bacteria indexes are built on 7,692 genomes from seven different bacterial species, then queried by 100K long ONT reads (Zymo). For HPRC experiments 1,188,163 long reads simulated from the CHM13 reference are used for the query. *Note that Movi 1 result on HPRC Release 2 is estimated as the run did not complete successfully due to high memory usage during index construction.

The next three points for Movi 2 correspond to sampled-*id* indexes with sampling rates of 50, 100, and 200. Higher sampling rates resulted in smaller index and slower query. Using different sampling rates, Movi 2 was able to use less memory than

SPUMONI in all three different datasets while remaining faster. In the Bacteria dataset, Movi 2’s memory usage became smaller than SPUMONI at sampling rate of 100 while being 4.86x faster. In HPRC Release 1, the sampled-*id* Movi 2 was smaller than both SPUMONI and ropebwt3 at all sampling rates we evaluated. For sampling rate of 50, Movi 2’s PML computation was 8.61x faster than SPUMONI ‘s PML computation. It was also 22.95x faster than ropebwt3’s SMEM finding mode, though SMEMs are more difficult result to compute, making these results hard to compare.

The HPRC Release 2 dataset had a high average run length that caused more length splitting in the sampled-*id* mode. For this reason, a higher sampling rate (200) was needed for Movi 2’s index to become smaller than SPUMONI ‘s. In that scenario Movi 2 was still 3.38x faster than SPUMONI. Overall, the results demonstrate that indexes based on move structure are able to achieve simultaneous speed/memory improvements compared to *r*-index based methods.

### 3.3. Parallelism and latency hiding

Latency hiding by prefetching is one of the features enabled in Movi because it is based on the move structure. Movi 1 and Movi 2 are able to concurrently process multiple reads (called strands) with a single thread to achieve latency hiding. Movi 2 further extends this by offering parallelization across multiple threads, in addition to concurrent read processing within each thread. Figure 4 shows how Movi 2’s performance scaled as the number of threads increased. For each thread count, we also evaluated the effect of prefetching. Our results showed that prefetching improved performance even under thread-level parallelization.

**Fig. 4:**
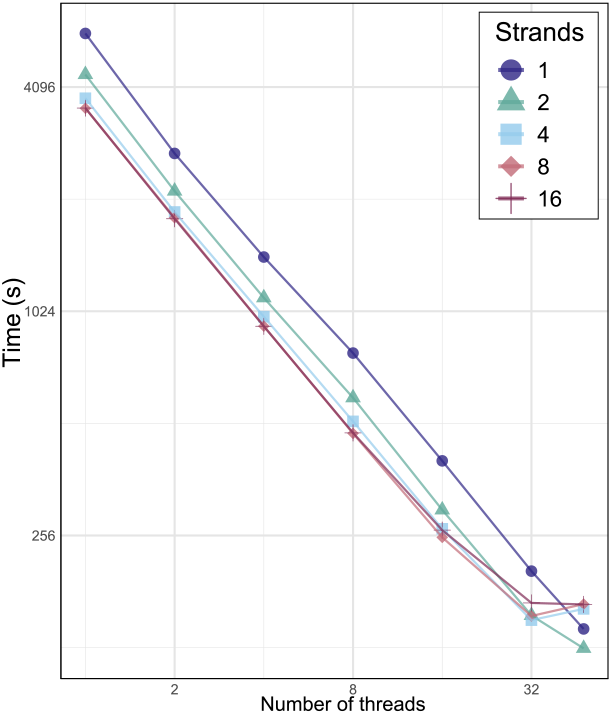
Movi 2’s performance with different number of threads and strands (for latency hiding). The number of concurrent reads processed per thread is referred to as strands.

We conducted experiments on a system with 48 hardware threads, scaling up to full utilization. We observed that when all threads are used, performance slows down, likely due to system saturation and competition for compute resources. In this case, prefetching can have an inverse effect, further intensifying resource contention.

## 4. Discussion

As pangenomes grow in size and utility, there is a growing need to improve the size and speed of pangenome indexes. The move structure is a compressed full-text index that has been shown to be drastically faster than others like the *r*-index. While this was thought to inevitably come at the price of larger data structure, here we showed that the move structure can be both smaller and faster than an *r*-index. We implemented these ideas in the Movi 2 software, and further showed that a range of these ideas yield a flexible trade-off space such that Movi 2 dominates either the *r*-index or the previous version (Movi) when configured appropriately.

A question for future work is whether the move structure, including the ideas proposed here, can be combined with minimizer digestion in order to improve the efficiency of sequence classification. This was shown to be possible using the r-index in the SPUMONI2 study (Ahmed et al., 2023). That study also showed that this has computational benefits, including (a) the addition of a layer of lossy compression (by way of minimizer digestion) on top of the lossless compression already achieved by the BWT, (b) a reduction in the number of iterations required in the matching inner loop, since the loop can proceed minimizer-by-minimizer rather than character-by-character. However, it is not yet known if the move structure can benefit similarly. An obstacle is that minimizer digestion in its most typical form will increase the size of the alphabet, since the alphabet is “promoted” from the A/C/G/T nucleotides to the larger set of possibly minimizer sequences. The SPUMONI2 study uses 4-character minimizers such that the minimizer alphabet has size 4^4^ = 256. Increasing the alphabet size introduces a particular challenge for storing thresholds in the current design of Movi 2. Each additional alphabet symbol requires its own threshold, since a new bit must be allocated to represent it. One way to address this is to store thresholds in two groups: one for characters with a threshold bit of 0 and another for those with a threshold bit of 1. We hypothesize that the number of distinct ways that characters will fall into these 0/1 groups will be limited, allowing them to be represented with an integer id that uses fewer than 256 bits. A further complication arises in repositioning, as it may now require scanning farther across nearby rows to locate a move row with the appropriate “character” (i.e. minimizer). Further investigation is required to determine if the move structure can maintain its efficiency advantage even for larger alphabets.

## 5. Author contributions statement

MZ and BL designed the method, with help from NKB. MZ wrote the software with assistance from NKB and TG. MZ performed the experiments. All authors contributed to the manuscript.

## 6. Acknowledgments

This work was supported by NIH grant R21HG013433 to BL.

## Supplementary Materials

### 1. Storing only the non-trivial thresholds

As an advanced version of Movi 1, we implemented a specific thresholds compression approach to store a threshold status instead of an offset for each character. A threshold-offset for each character requires *O*(log_2_(*ℓ*_*max*_)) bits (2 bytes in Movi), whereas a threshold-status requires only 2 bits. This is because the status needs to express three main possibilities (assuming the length of the move row is *ℓ*):

1. The threshold is at the start of the move row (0).
2. The threshold is at the end of the move row (*ℓ*).
3. a non-trivial offset inside the move row (between 0 and *ℓ*).

The first two cases do not require any additional information to retrieve the threshold offset. In case (1), the threshold offset is 0 and in case (2), it is set to the length of the the move row. For case (3), the explicit offset of the non-trivial threshold is stored in a dedicated field which requires *O*(log_2_(*ℓ*_*max*_)). However, this is the only column requiring *O*(log_2_(*ℓ*_*max*_)) bits, which is an improvement over storing Σ − 1 columns with the same number of bits. It is also possible, though rare in our experiments, for multiple non-trivial thresholds to exist in a row. This can be identified using the 4^th^ code of the threshold status. To handle such cases, we store an overflow table that explicitly stores the thresholds for rows with more than one non-trivial threshold. In practice, we find that this table is small compared to the main Movi table. This approach was implemented in a later release of Movi 1, reducing the total size of each move row from 16 bytes to 12 bytes.

When a trivial threshold is represented by a threshold status rather than full offset, the most significant part of the threshold data structure in Movi becomes the non-trivial thresholds.

### 2. The blocked index

### 3. Computing the *id* in the sampled-*id* mode

#### Algorithm 1

Computing the *id* for move row *q* using the *S* table. *M* is the Move table, *s* is the sampling rate.

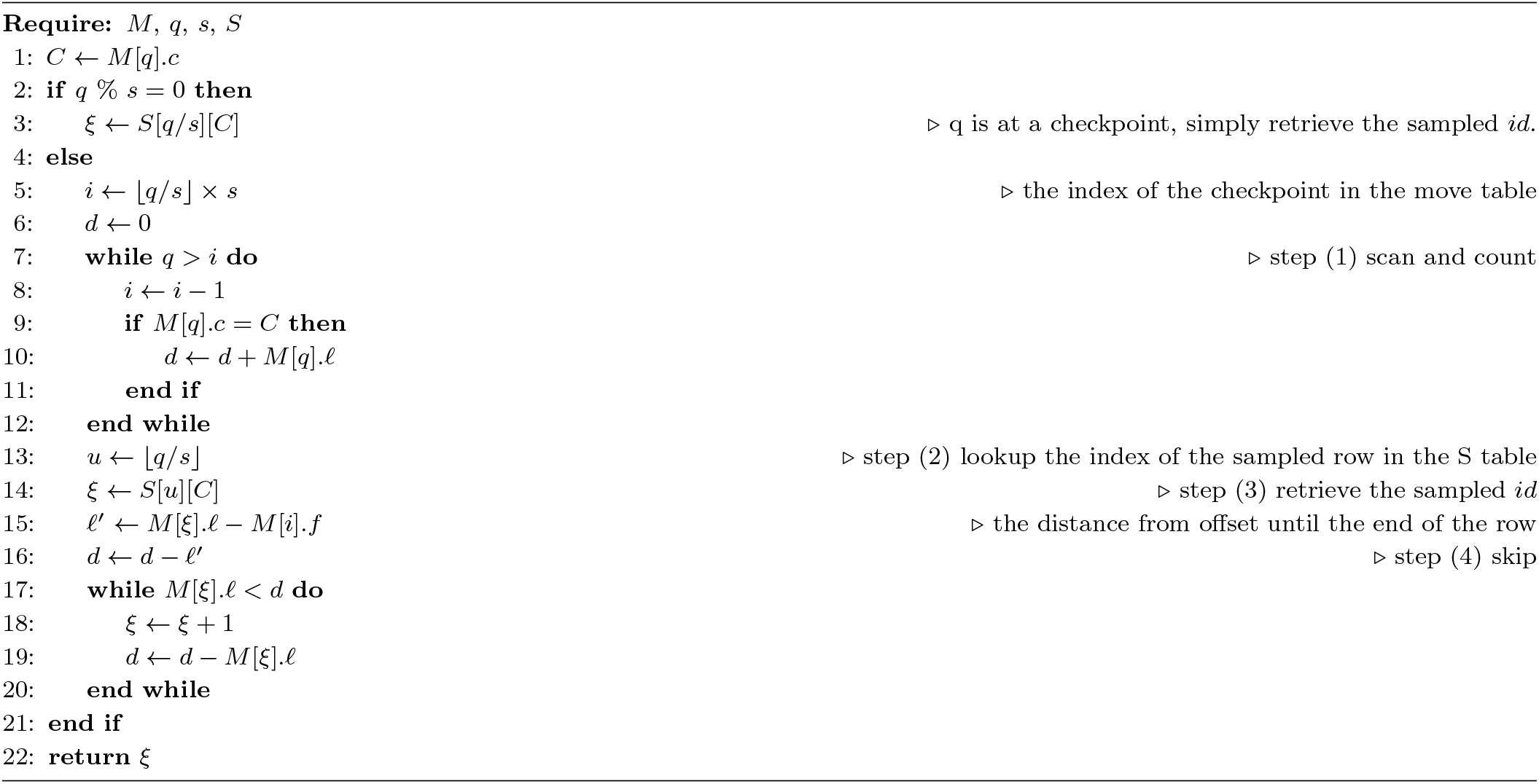

## Notes

https://github.com/mohsenzakeri/Movi

### Competing Interest Statement

The authors have declared no competing interest.

### Summary of Updates

I only needed to update the funding information in the web page (not inside the paper). The correct information is "NIH R21HG013433". Thank you!

